# The mitochondrial Rho-GTPase, Miro, is resident at peroxisomes and regulates peroxisomal trafficking and morphology

**DOI:** 10.1101/241208

**Authors:** Christian Covill-Cooke, Guillermo López-Doménech, Nicol Birsa, Josef T. Kittler

## Abstract

Peroxisomes are essential for a number of cellular functions, including reactive oxygen species metabolism, fatty acid β-oxidation and lipid biosynthesis. To ensure optimal functionality of peroxisome-dependent processes throughout the cell they must be trafficked; however, peroxisomal transport remains poorly characterised. Here we show that Miro1 and Miro2, outer mitochondrial membrane proteins essential for mitochondrial trafficking, are also localised to peroxisomes. Peroxisomal localisation of Miro1 is negatively regulated by its first GTPase domain and is mediated by an interaction through its transmembrane domain with the peroxisomal-membrane protein chaperone, Pex19. By using Miro1/2 double knockout mouse embryonic fibroblasts (MEFs) we find that the loss of Miro1/2 leads to a significant reduction in short-range microtubule-independent peroxisomal motility. Additionally, Miro regulates peroxisomal size and morphology. Our results contribute to the fundamental understanding of peroxisomal trafficking and morphology, supporting a complex crosstalk between peroxisomal and mitochondrial biology.

## Introduction

Peroxisomes are single-membrane bound organelles that are required for a wide range of essential metabolic pathways. As sites of both the production and clearance of reactive oxygen species (ROS) as well as the biosynthesis of specific lipids (e.g. plasmalogens), peroxisomes are critical for cellular health. This is emphasised by loss-of-function mutations of key genes in peroxisomal biogenesis *(PEX* genes) leading to Zellweger spectrum disorders (Klouwer et al., 2015). As peroxisomes have a role in metabolism, they are known to respond to environmental cues by altering their size, number and distribution to ensure optimal functionality (Smith and Aitchison, 2013).

Peroxisomes can be generated *de novo* by the combination of pre-peroxisomal vesicles from the endoplasmic reticulum and mitochondria (Agrawal and Subramani, 2016; Kim et al., 2006; Sugiura et al., 2017). However, more often they are generated from pre-existing peroxisomes through a process that requires peroxisomal elongation driven by Pex11β and subsequent peroxisomal fission (Ebberink et al., 2012; Koch and Brocard, 2012). Strikingly, peroxisomal fission requires overlapping machinery with mitochondrial fission, with hFis1 and Mff being localised to peroxisomes for the recruitment of Drp1 (Gandre-Babbe and van der Bliek, 2008; Koch et al., 2005, 2003). The peroxisomal targeting of hFis and Mff is proposed to occur from the cytosol by the membrane protein chaperone, Pex19, suggesting an axis whereby these proteins can be either targeted to the mitochondria or peroxisomes (Delille and Schrader, 2008).

As peroxisomes are involved in a diverse range of metabolic functions and the fact that they interact with several organelles (Shai et al., 2016), peroxisomes must be trafficked throughout the cell. The importance of this has been emphasised by *SPAST* mutant (associated with hereditary spastic paraplegia) cells exhibiting reduced peroxisomal trafficking and subsequently defects in distribution, resulting in impaired handling of ROS (Abrahamsen et al., 2013; Wali et al., 2016). The current paradigm of peroxisomal trafficking in mammalian cells is that ~5-15% of peroxisomes undergo long-range microtubule-dependent trafficking, with the rest exhibiting shorter-range displacements, often referred to as oscillatory motility. Despite the importance of peroxisomal trafficking, the mechanisms by which trafficking occurs are poorly understood (Bharti et al., 2011; Huber et al., 1997, 1999; Rapp et al., 1996; Schrader et al., 2000; Wiemer et al., 1997).

The mitochondrial Rho-GTPases, Miro1 and Miro2, are outer mitochondrial membrane proteins critical for mitochondrial trafficking (Birsa et al., 2013; López-Doménech et al., 2016, 2018; Macaskill et al., 2009; Wang and Schwarz, 2009). Structurally, both Miro paralogues exhibit a large, cytoplasm-facing N-terminal domain with two calcium binding EF-hand domains flanked by a GTPase domain on each side (Fransson et al., 2003; Klosowiak et al., 2013). Here we show that Miro1 and Miro2 are not strictly localised to mitochondria but are also localised to peroxisomes. Moreover, this peroxisomal localisation of Miro is regulated through its first GTPase domain. We find that, through its transmembrane domain, Miro1 can interact with Pex19 suggesting targeting to the peroxisome from the cytosol. Utilising Miro1/2 knockout (DKO) MEFs we find that Miro regulates microtubule-independent peroxisomal motility and peroxisomal size.

## Results

Whilst imaging the subcellular localisation of GFP tagged Miro1 (^GFP^Miro1) and Miro2 (^GFP^Miro2) in mouse embryonic fibroblasts (MEFs) we noticed that alongside their well-documented mitochondrial localisation (Fransson et al., 2003), Miro1 and Miro2 fluorescent signal could also be observed on small vesicular structures. Colocalisation of these structures with catalase staining confirmed that these structures were in fact peroxisomes (Figure 1A). To measure the extent of this peroxisomal localisation, thresholded GFP signal on catalase positive but Tom20 negative structures was quantified (see Material and Methods). Both ^GFP^Miro1 and ^GFP^Miro2 showed a significant enrichment of peroxisomal localisation over ^GFP^Tom70(1-70) (a GFP fusion protein of the mitochondria-targeting sequence of Tom70 (amino acids 1-70)), highlighting a specific localisation of Miro to the peroxisomes and not simply a mislocalisation of outer mitochondrial membrane (OMM) proteins (Figure 1A-B). Furthermore, the peroxisomal localisation of Miro was observed with a ^myc^Miro1 construct and found to also occur in Cos7 cells (Supplementary Figure 1).

hFis 1 exhibits both a mitochondrial and peroxisomal localisation and can induce mitochondrial and peroxisomal fission (Koch et al., 2005). The peroxisomal localisation of hFis1 is dependent on the ability of its C-terminal transmembrane domain to bind to Pex19; a chaperone that targets newly synthesised peroxisomal membrane proteins from cytosolic ribosomes to peroxisomes. We tested whether Miro1, which also contains a C-terminal transmembrane domain, can similarly interact with Pex19 and whether this interaction is dependent on its transmembrane domain. To achieve this we expressed ^myc^Pex19 and both full-length Miro1 and Miro1 lacking the transmembrane domain in Cos7 cells. By pulling down Pex19 we observed robust co-immunoprecipitation of Miro1 and Pex19 which was completely abolished upon deletion of the transmembrane domain of Miro1 (Figure 1C). Therefore, we conclude that both Miro1 and Miro2 can localise to peroxisomes and that the transmembrane domain of Miro1 is critical for its interaction with Pex19.

Being anchored in the OMM by their C-terminus, both Miro1 and Miro2 exhibit a large cytoplasm-facing N-terminus. Structurally, the N-terminal part of the proteins includes two EF hand domains flanked by a GTPase domain on either side. To characterise the importance of these domains in the peroxisomal localisation of Miro, we generated truncation constructs of Miro1 (Figure 2A) and expressed them in Miro1 / Miro2 double knockout (DKO) MEFs to prevent the influence of any endogenous Miro. Strikingly Miro1 lacking the first GTPase domain (^gfp^EF1-EF2-GTP2) exhibited a dramatic increase in peroxisomal localisation in comparison to either full-length Miro1 or GTPase2 domain (^gfp^GTP2) alone (Figure 2B-C). In contrast, C-terminally anchored GTPasel domain (^gfp^GTP1), which was still effectively targeted to mitochondria did not show any enrichment on peroxisomes compared to ^GFP^Tom70(1-70). Given the importance of the first GTPase domain to the localisation of Miro1, we tested whether it influences the binding of Miro1 to Pex19. Cos7 cells were transfected with both the ^GFP^Miro1 truncation constructs (Figure 2A) and ^myc^Pex19. Following pulldown of GFP, ^myc^Pex19 was found to co-immunoprecipitate with the full-length ^GFP^Miro1, ^gfp^EF1-EF2-GTP2 and ^gfp^GTP2 forms of Miro1. Interestingly, binding of ^myc^Pex19 and ^gfp^GTP1 occurred to a much lesser extent (Figure 2D) suggesting that the first GTPase domain may negatively regulate the ability of Pex19 to bind to the transmembrane domain of Miro1. This, along with the observation of enhanced peroxisomal localisation of ^gfp^EF1-EF2-GTP2 (lacking GTPase domain 1; Figure 2B and C), suggests GTPase domain 1 plays an important regulatory role in the peroxisomal targeting of Miro1.

Following the identification of key features required for the peroxisomal localisation of Miro, we next sought to better understand the function of Miro at peroxisomes. Accounting for ~10% of peroxisomal transport, long-range peroxisomal trafficking (known as saltatory trafficking from hereon) is well characterised as being microtubule-dependent (Huber et al., 1997; Rapp et al., 1996; Schrader et al., 2000, 1996; Wiemer et al., 1997) and thought to occur in part by the kinesin-1 family of motors (Kulic et al., 2008). The rest of peroxisomal transport (~90%) occurs by shorter-range oscillatory motility (Bharti et al., 2011; Huber et al., 1997; Rapp et al., 1996; Wiemer et al., 1997). Miro has been extensively documented to be critical for bi-directional, microtubule-dependent trafficking of mitochondria in a wide variety of species and cell types (Birsa et al., 2013; Schwarz, 2013; Stephen et al., 2015; Vaccaro et al., 2017; López-Doménech et al., 2018) suggesting that the peroxisomal pool of Miro could also be important for microtubule-dependent peroxisomal transport. To observe the Miro-dependence of saltatory peroxisomal trafficking, pxDsRed (DsRed2 localised to the peroxisomal lumen by a PTS1; a peroxisomal lumen targeting signal) was transfected into wild type (WT) and Miro DKO MEFs and imaged at one frame every 1.5 seconds for two minutes (Supplementary movie 1 (WT) and 2 (DKO)). Surprisingly, quantification of saltatory events over two-minute movies by blind scoring showed no difference in this behaviour in DKO MEFs compared to wild type (WT: 8.50 ± 0.66, DKO: 7.63 ± 1.44 events per cell, *p*=0.5710; Figure 3A; Supplementary movie 3 (WT) and 4 (DKO)). Depolymerisation of microtubules by vinblastine abolished long-ranged saltatory trafficking, as reported previously (Supplementary figure 2; Supplementary movie 5 (WT) and 6 (DKO)) (Bharti et al., 2011; Huber et al., 1997; Rapp et al., 1996; Salogiannis et al., 2016; Schrader et al., 2000, 1996; Wiemer et al., 1997). Therefore, the genetic loss of both Miro1 and Miro2 does not have an effect on long-ranged microtubule-dependent trafficking of peroxisomes.

Alongside its role in microtubule-dependent trafficking of mitochondria, the loss of Miro1 has also been shown to dramatically affect the positioning of mitochondria, with mitochondria becoming more perinuclear in distribution (Kanfer et al., 2015; López-Doménech et al., 2016, 2018; Nguyen et al., 2014). To quantify whether there is a difference in the distribution of peroxisomes between WT and DKO MEFs, a Sholl-like quantification method was applied (as published previously (López-Doménech et al., 2016, 2018). Briefly, MEFs were seeded on fibronectin micropatterns to standardise cell morphology, then fixed and stained for catalase. Peroxisome distribution was then measured by concentric circles being drawn from the centre of the cell at 1 µm intervals and the inter-circle catalase signal being quantified (Supplementary Figure 3A) (López-Doménech et al., 2016, 2018). As seen in the normalised cumulative distribution of catalase signal, knockout or overexpression of Miro does not cause an alteration in the overall distribution of peroxisomes (Figure 3B-C and Supplementary figure 4D). Quantification of the distance at where 50% (Perox^50^) and 95% (Perox^95^) of peroxisomes are situated, further supports this conclusion (Figure 3D; Supplementary Figure 3B & 4A-D). The distribution of mitochondria upon loss Miro1/2 on the other hand was dramatically affected, in agreement with previous work (Figure 3B; Supplementary Figure 3C-D) (López-Doménech et al., 2018). As a result, despite the role of Miro in microtubule-dependent mitochondrial trafficking and mitochondrial distribution, Miro does not affect saltatory peroxisomal trafficking or proximo-distal peroxisomal distribution.

Approximately 90% of peroxisomal trafficking occurs by shorter-range transport, often referred to as oscillatory motility. In contrast to saltatory peroxisomal trafficking the mechanism by which shorter-range oscillatory motility is elicited is not well-defined, with conflicting data stating that it is either caused by ATP-dependent random motion in the cytosol or by association with the actin cytoskeleton (Bharti et al., 2011; Huber et al., 1997; Wiemer et al., 1997). As Miro is not required for saltatory peroxisomal trafficking we tested whether or not it might have a role in shorter-range displacements. To test this, individual peroxisomes were tracked and their oscillatory behaviour measured through displacement of the peroxisomes over time. Interestingly, we found that loss of Miro leads to a ~30% reduction in the median net displacement of peroxisomes (Figure 4A-B); meaning that Miro is required for the short-ranged motion of peroxisomes. Depolymerisation of microtubules had no effect on median net displacement, further supporting the role of Miro in microtubule-independent trafficking of peroxisomes (Supplementary figure 2C). Consequently, Miro is a novel regulator of the oscillatory behaviour of peroxisomes in a mechanism independent of microtubules.

Coupled to its role in mitochondrial trafficking, overexpression and loss of Miro has been shown to lead to long reticulated and short rounded mitochondria, respectively (López-Doménech et al., 2018; Ding et al., 2016; MacAskill et al., 2009; Russo et al., 2009; Saotome et al., 2008). As a result, the possibility of a role for Miro in peroxisomal morphology was examined. Peroxisomes are known to adopt either a vesicular or tubular morphology with an average diameter ranging between 0.1 μm and 1 μm, depending on cell type and environmental cues (Smith and Aitchison, 2013). By staining peroxisomes in WT and DKO MEFs with a catalase antibody we observed that peroxisomes in the DKO MEFS were smaller and less tubular; as depicted by a decrease in average catalase puncta area (Figure 4C-D). In contrast, overexpression of Miro1 in WT MEFs led to an increase in average peroxisome size (Figure 4E-F). Thus, the expression levels of Miro can regulate peroxisomal size.

All in all, we have identified novel non-mitochondrial roles for Miro in the regulation of peroxisomal motility and size. Through an interaction with Pex19, Miro localises to peroxisomes by a mechanism dependent on its transmembrane domain and signalling through the first GTPase domain. Interestingly, Miro does not regulate microtubule-dependent trafficking of peroxisome, nor their basal distribution, but rather is important for shorter-range movements and oscillatory motility. Not only do these results provide a mechanism for short-ranged peroxisomal motility, but also highlight Miro as a regulatory link between peroxisomal trafficking and size.

## Discussion

One question that arises from the dual localisation of Miro to mitochondria and peroxisomes is how are the relative pools of Miro on each organelle achieved? Recently, Okumoto *et al*. (2018) provided evidence that alternative splicing of exon 19 in Miro1, at a site near to the TM domain, leads to its enhanced peroxisomal localisation. Interestingly, from their *in vitro* binding assays they propose that Miro1 is targeted to peroxisomes by exon 19 sequences interacting with Pex19. Our data, however, show that Miro proteins lacking the exon 19 splice cassette including Miro2 and the most common Miro1 variant (variant 1) can also be found targeted to peroxisomes, in agreement with a recent analysis of C-terminally anchored proteins (Costello et al., 2017). Additionally, we show that the transmembrane domain of Miro is necessary for the interaction with Pex19 in cells, and by extension the localisation of Miro to peroxisomes. Indeed, Pex19 has been shown to bind the transmembrane domain of its targets (Delille and Schrader, 2008; Halbach et al., 2006). Moreover, the subcellular localisation of C-terminally anchored proteins is well documented as being dependent on the biochemical properties of the transmembrane domain and C-terminal amino acids (Borgese and Fasana, 2011; Borgese et al., 2007; Costello et al., 2017; Halbach et al., 2006). As a result, we propose a model whereby the Miro transmembrane is required for Pex19 binding and peroxisomal localisation of Miro. Therefore, other features such as the first GTPase domain and sequences within exon 19 may act as important sites for regulatory factors to bind and modulate the extent of the peroxisomal pool of Miro. In agreement with this, we also show that loss of GTPase domain 1 can enhance peroxisomal targeting of Miro1 in the absence of exon 19 sequences. Consequently, the ability to control the extent of the mitochondrial and peroxisomal localisation of Miro may be an important regulatory axis.

Miro plays an important role in establishing a properly distributed mitochondrial network in many cell types through long-range microtubule-dependent trafficking (Kanfer et al., 2015; López-Doménech et al., 2016, 2018; Nguyen et al., 2014). Indeed, we have shown that genetically knocking out all Miro1 and Miro2 in MEFs halts long-range mitochondrial trafficking leading to a dramatic perinuclear collapse of the mitochondrial network (López-Doménech et al., 2018). A recent study also reported that a peroxisomally localised splice variant of Miro1 modulates long-ranged trafficking and subsequent redistribution of peroxisomes in HeLa cells (Okumoto et al., 2017). Here, using a combination of micropattern based cell standardisation and quantitative organelle distribution analysis, we now show that compared to the marked collapse of mitochondria upon Miro1/2 DKO, steady state peroxisomal distribution remains unaffected. In addition, while we have previously shown that knocking out all Miro leads to a dramatic reduction in directed microtubule-dependent transport of mitochondria we could find no evidence for the involvement of Miro in long-range microtubule-dependent trafficking of peroxisomes. Thus, unlike for mitochondria, the primary role of Miro on peroxisomes does not appear to be to mediate their correct distribution throughout the cell.

In contrast, we find that Miro is an important regulator of shorter-range oscillatory motility; a type of trafficking that makes up ~90% of all peroxisomal movement (Huber et al., 1997; Schrader et al., 1996; Wiemer et al., 1997). With roles for peroxisomes in many aspects of metabolism, oscillatory motility may be important for peroxisomal surveillance of the cell and in maintaining the ability to respond to dynamic changes in intracellular microenvironments. In fact, despite making up a tiny fraction of the cell, peroxisomes can explore up to ~30% of the cytoplasmic volume over a 15 minute period (Valm et al., 2017). Furthermore, their interactions with other organelles can respond to changes in cellular metabolism (Valm et al., 2017). Unlike long-range saltatory peroxisomal trafficking, oscillatory motility has often been dismissed as simply Brownian-like movement (Huber et al., 1997; Rapp et al., 1996; Wiemer et al., 1997). Not only is Miro the first described peroxisomally localised regulator of short-range oscillatory trafficking, our work also provides support for an active mechanism controlling shorter-range peroxisomal dynamics (Bharti et al., 2011). Mechanistically we show that microtubules are not required for this oscillatory motility, in agreement with previous work (Bharti et al., 2011; Huber et al., 1999; Kulic et al., 2008; Rapp et al., 1996; Schrader et al., 2000; Wiemer et al., 1997). Given the previously described roles for the actino-myosin cytoskeleton in peroxisomal trafficking (Bharti et al., 2011; Lin et al., 2016; Schollenberger et al., 2010), it might be that the role of Miro in shorter-range oscillatory motility is via an actin-dependent mechanism. Indeed, we recently demonstrated a role for Miro in conjunction with actin and myosin-19 in short-range mitochondrial displacements (López-Doménech et al., 2018). Consequently, Miro may play a similar role, stabilising myosin-19 or another myosin on peroxisomes to facilitate their actin-dependent movement or anchoring (Lin et al., 2016; Schollenberger et al., 2010; López-Doménech et al., 2018).

Through addition of membrane from the endoplasmic reticulum along with peroxisomal fission, peroxisome morphology is known to be highly dynamic and able to respond to environmental signals such as variations in metabolism (Smith and Aitchison, 2013). Alongside controlling peroxisomal trafficking, we show that Miro regulates peroxisomal size. The ability to regulate peroxisome number and morphology is not only important in controlling the abundance of peroxisomes within the cell, but may also affect enzyme activity of individual peroxisomes (Kiel et al., 2005). Interestingly, peroxisome-ER contacts were recently demonstrated to be an important determinant of peroxisome growth (Hua et al., 2017). As Miro was previously demonstrated to play a role in coupling mitochondria to the ER (Lee et al., 2016), it may be important in mediating peroxisome-ER contacts, subsequently regulating peroxisomal dynamics.

At mitochondria, Miro has a multifaceted role covering several aspects of mitochondrial dynamics, function and turnover and therefore is proposed to be important in maintaining cellular health (Birsa et al., 2014, 2013; Covill-Cooke et al., 2017; Devine et al., 2016; Misko et al., 2010; Weihofen et al., 2009). As our data uncovers a role for Miro in peroxisomal dynamics, it is important to consider the significance of this protein being localised at both mitochondria and peroxisomes. Mitochondria and peroxisomes share several key functions and have been suggested to be evolutionarily related (Andrade-Navarro et al., 2009). For example, both are sites of fatty acid (β-oxidation, lipid synthesis and have a role in ROS metabolism. As a result, the ratio of mitochondrial to peroxisomal localisation of Miro could act as a means to coordinate the function of both organelles in a dynamic cellular environment.

## Materials and Methods

### Antibodies and reagents

DNA constructs: ^GFP^Miro1, ^GFP^Miro2 and ^myc^Miro1 have been described previously (Fransson et al., 2003); ^GFP^Tom70(1-70), amino acids 1-70 of human Tom70 cloned into pEGFP-N1; Miro1 truncation constructs were cloned from ^GFP^Miro1; ^myc^Miro1△TM cloned from ^myc^Miro1, pxDsRed from Addgene (#54503), ^myc^Pex19 mouse Pex19 (NM_023041) cloned into pRK5-myc vector. Primary antibodies for immunofluorescence (IF) and western blotting (WB): rabbit anti-Tom20 (Santa Cruz sc-11415, IF 1:500), mouse anti-Catalase (Abcam ab110292, IF 1:500), rabbit anti-Pex14 (Atlas HPA04386, IF 1:500), mouse anti-myc supernatant (purified in house from 9E10 hybridoma cell line, WB 1:100), rat anti-GFP (nacalai tesque 04404-84, IF 1:2,000), rabbit anti-Pex19 (Abcam ab137072, WB 1:1000) and rabbit anti-GFP (Santa Cruz sc-8334, WB 1:100). Fluorescent secondary antibodies (all from Thermo Fisher Scientific, 1:1,000): Donkey anti-rat Alexa Fluor-488 (A21208), goat anti-rabbit Alexa Fluor-555 (A21430) and donkey anti-mouse Alexa Fluor-647 (A31571).

WT and DKO MEFs were characterised previously (López-Doménech et al., 2018). For peroxisomal distribution analysis, MEFs were seeded onto large-Y-shaped-fibronectin-micropatterned coverslips (CYTOO 10-012-00-18) at a density of 15,000 to 20,000 cells/cm^2^. Cells were then left to attach for three hours and then fixed for 10 minutes with 4% paraformaldehyde (PFA). Immunocytochemistry was then carried out as described below.

### Co-immunoprecipitation and western blot analyses

Cells were lysed in buffer containing 50 mM Tris-HCl pH 7.5, 0.5% Triton X-100, 150 mM NaCl, 1 mM EDTA, 1 mM PMSF and protease inhibitor cocktail for 45 minutes at 4°C with rotation. Lysates were then centrifuged at 14,000 rpm for 15 minutes and the supernatant collected for inputs and subsequent immunoprecipitation. GFP-tagged proteins were pulled down with GFP-trap agarose beads (Chromotek, gta-10) for 2 hours. Beads were then washed three times with the lysis buffer.

Samples were run on SDS-PAGE gel and transferred onto nitrocellulose membrane. Membranes were blocked with 4% (w/v) milk in PBS with 0.1% Tween20 (PBST). Primary antibodies were incubated overnight at 4°C, washed three times with PBST and incubated with the secondary for 45 minutes at room temperature. Following three washes with PBST the membrane was developed by exposure to ECL substrate (Millipore, WBLUR0500) and imaged on the ImageQuant LAS4000 mini (GE Healthcare).

### Fixed imaging

Cells were fixed with 4% PFA for 10 minutes and blocked for 30 minutes with 10% horse serum, 5 mg/ml bovine serum albumin and 0.2% Triton X-100 in PBS. Samples were stained with primary and secondary antibodies for one hour each and imaged on a Zeiss LSM700 confocal using a 63x oil objective (NA=1.4).

### Live imaging

Live imaging of pxDsRed in WT and DKO MEFs was carried out at 37°C whilst perfusing a solution of 10 mM glucose, 10 mM HEPES, 125 mM NaCl, 5 mM KCl, 2 mM CaCl and 1 mM MgCl at pH 7.4 by addition of NaOH, onto the coverslips. A 60x water objective on an Olympus BX60M microscope with an Andor iXon camera was used to acquire images. MicroManager software was utilised to control the microscope set-up. PxDsRed was excited with an ET548/10x filter. To depolymerise microtubules Vinblastine was added at 1 μm for 1 hour prior to imaging.

### Image analysis

Quantification of the extent of peroxisome localisation was carried out in ImageJ. Using Tom20 signal as a mitochondrial mask, all GFP signal that overlapped with the mitochondria was removed. GFP signal that then colocalised with catalase positive structures was thresholded (kept constant for all conditions) and then the area measured using Analyze Particles. Peroxisomal morphology was measured by quantification of thresholded catalase signal in ImageJ. Both area and Feret's diameter were measured. Live trafficking of pxDsRed signal was quantified using TrackMate (Tinevez et al., 2017). Only tracks that last lasted more than half the movie were used for analysis to prevent peroxisomes occurring more than once in the dataset.

### Statistical analysis

Graphpad prism was used to statistically analyse data. For comparisons between two conditions a two-tailed Student's t-test was used. For multiple comparisons either a one-way ANOVA with a Tukey post-hoc test or Kruskall-Wallis with a post-hoc Dunn's correction was used as stated in the figure legends. Graphed data is presented as mean plus or minus S.E.M.

## Acknowledgements

We thank all members of the Kittler lab for helpful discussions and comments. This work was supported by a European Research Council grant (Fuelling Synapses) and Lister Institute of Preventive Medicine Award to J.T.K. C.C-C is a recipient of a studentship from the Medical Research Council (1368635).

## Author contributions

C.C-C, G.L-D and J.T.K designed experiments. C.C-C. collected and analysed the results. G.L-D. generated the WT and DKO MEFs and cloned the ^GFP^Tom70(1-70) construct. N.B. cloned the Miro1 truncation constructs. C.C-C. and J.T.K. wrote the manuscript.

---

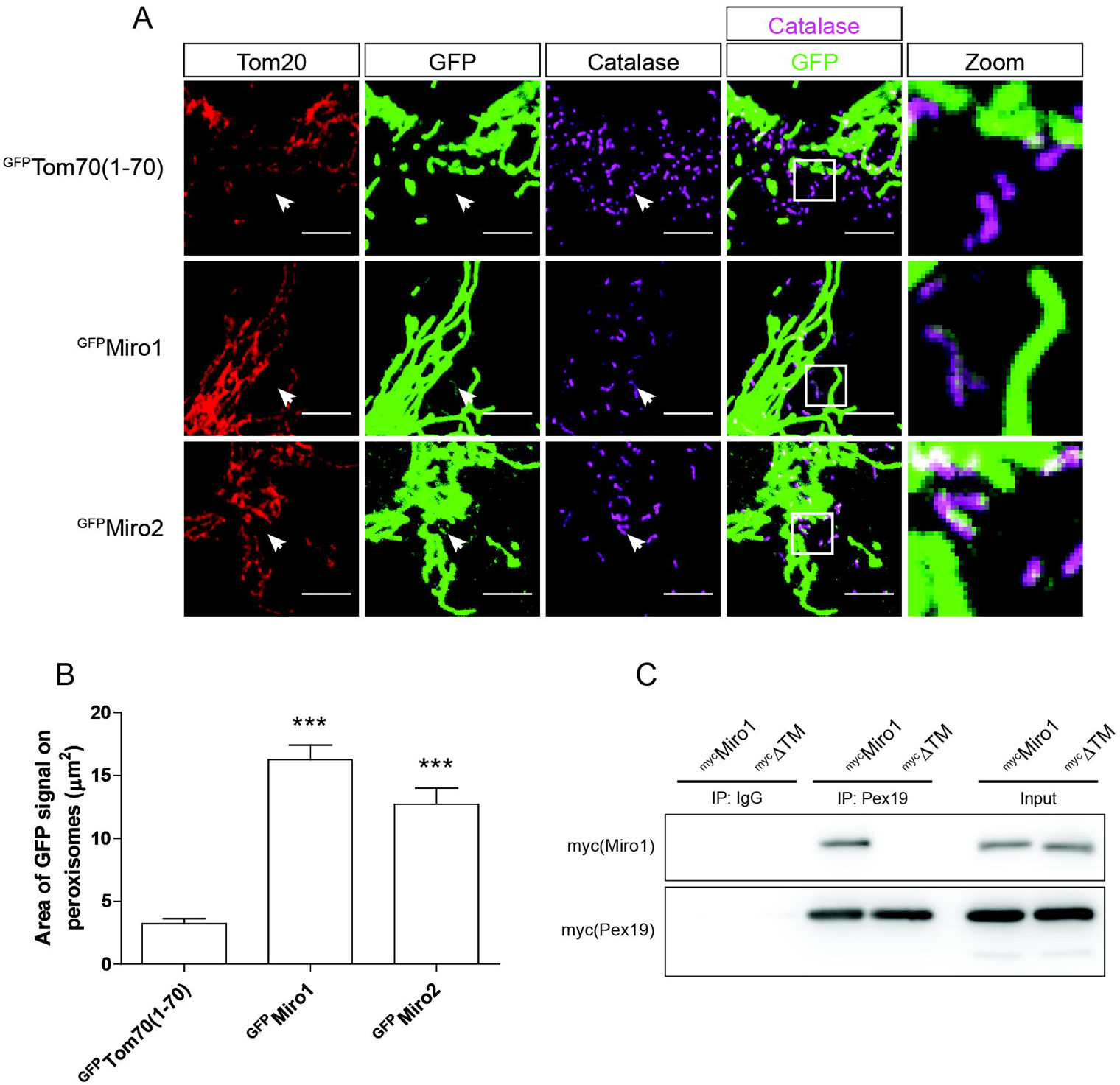

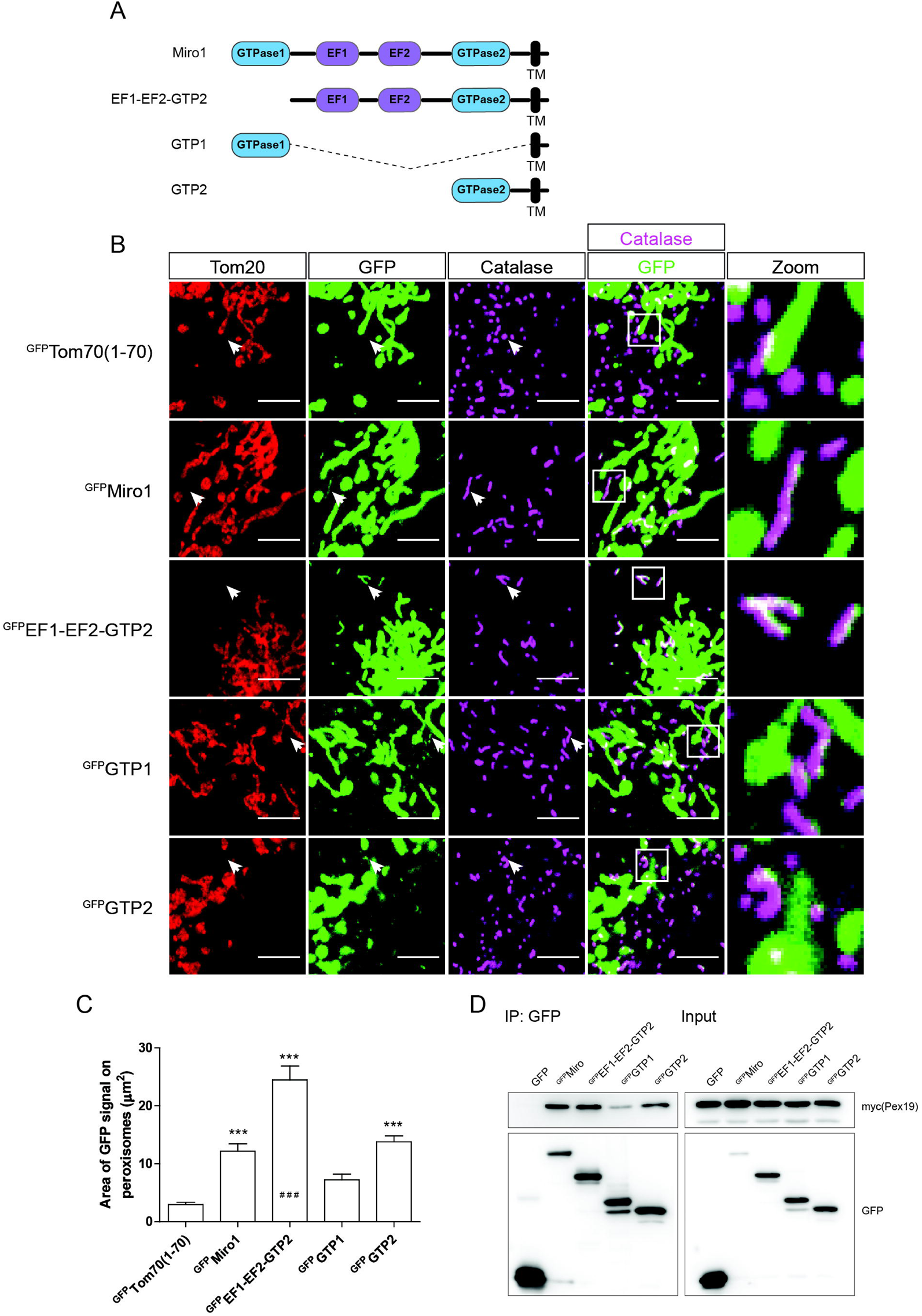

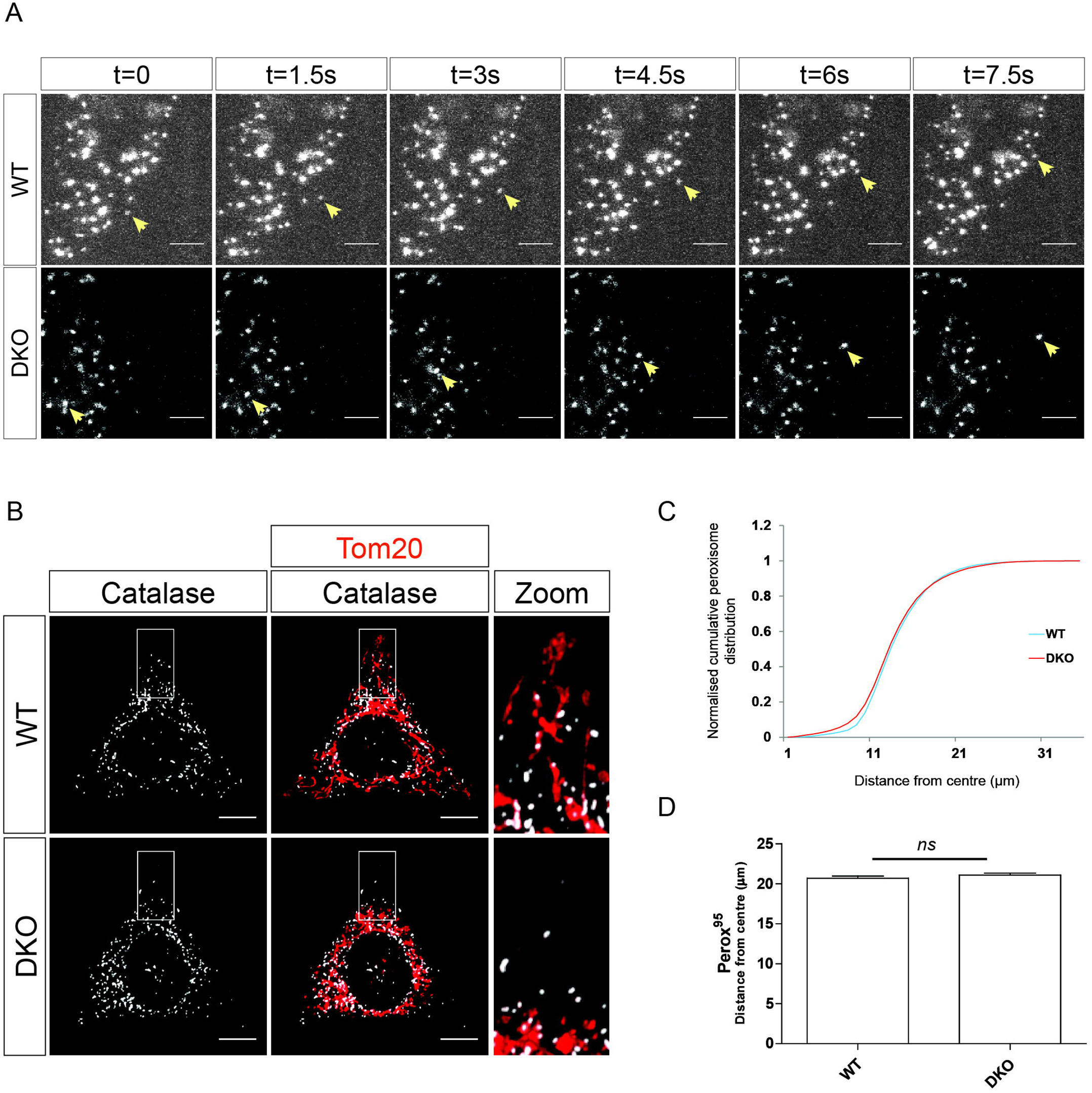

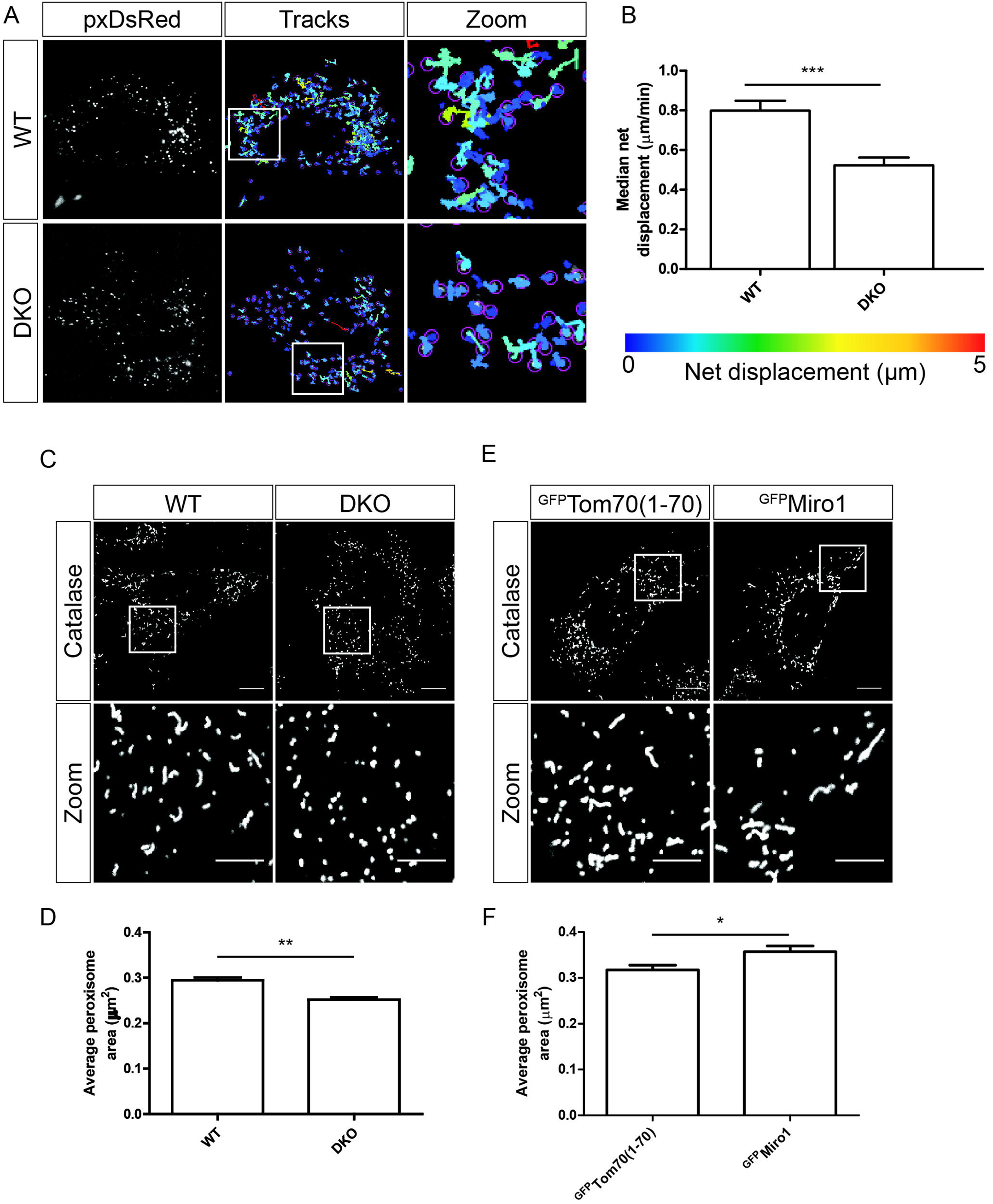

